# Criticality and universality in neuronal cultures during ‘up’ and ‘down’ states

**DOI:** 10.1101/2024.01.10.575061

**Authors:** Mohammad Yaghoubi, Javier G. Orlandi, Michael A. Colicos, Jörn Davidsen

## Abstract

The brain can be seen as a self-organized dynamical system that optimizes information processing and storage capabilities. This is supported by studies across scales, from small neuronal assemblies to the whole brain, where neuronal activity exhibits features typically associated with phase transitions in statistical physics. Such a critical state is characterized by the emergence of scale-free statistics as captured, for example, by the sizes and durations of activity avalanches corresponding to a cascading process of information flow. Another phenomenon observed during sleep, under anesthesia, and in *in vitro* cultures, is that cortical and hippocampal neuronal networks alternate between “up” and “down” states characterized by very distinct firing rates. Previous theoretical work has been able to relate these two concepts and proposed that only up states are critical whereas down states are subcritical, also indicating that the brain spontaneously transitions between the two. Using high-speed high-resolution calcium imaging recordings of neuronal cultures, we test this hypothesis here by analyzing the neuronal avalanche statistics in populations of thousands of neurons during “up” and “down” states separately. We find that both “up” and “down” states can exhibit scale-free behavior when taking into account their intrinsic time scales. In particular, the statistical signature of “down” states is indistinguishable from those observed previously in cultures without “up” states. We show that such behavior can not be explained by network models of non-conservative leaky integrate-and-fire neurons with short-term synaptic depression, even when realistic noise levels, spatial network embeddings, and heterogeneous populations are taken into account, which instead exhibits behavior consistent with previous theoretical models. Similar differences were also observed when taking into consideration finite-size scaling effects, suggesting that the intrinsic dynamics and self-organization mechanisms of these cultures might be more complex than previously thought. In particular, our findings point to the existence of different mechanisms of neuronal communication, with different time scales, acting during either high-activity or low-activity states, potentially requiring different plasticity mechanisms.

**Author summary:** Up and down states, where populations of neurons transition between periods of high and low-frequency activity, are ubiquitous in the brain. They are present during development, sleep, and anesthesia, and have been associated with memory consolidation and the regulation of homeostatic processes. Using large-scale high-speed calcium imaging recordings of neuronal cultures, we show that self-similar behavior can appear during both up and down states, but with different characteristic timescales. Detailed simulations of neuronal cultures are only able to capture the statistics during up states, suggesting that a different mechanism might be governing the dynamics of the down states. The presence of scale-free statistics with switching time scales points to novel self-organization mechanisms in neuronal systems.

## Introduction

The description of neuronal dynamics within the framework of critical phenomena has become commonplace amongst physicists in the last decade or so (1–3). The first signatures of criticality in the brain were depicted as neuronal avalanches in organotypic cortical cultures (4, 5), where sequences of high-frequency neuronal activations across a small population could be described by scale-free statistics of their sizes and durations distributions. Nowadays, similar critical signatures have been observed across many systems and preparations: from power-law statistics of correlations in whole-brain recordings (6, 7) to neuronal avalanches in slices (8), dissociated cultures (9, 10), and *in vivo* (11– 16). They indicate the emergence of complex spatiotemporal dynamics with statistics compatible with a system being in the neighborhood of a critical point, in particular, near a second-order phase transition, or of a critical region (17).

From a statistical physics point of view, neuronal avalanches are interpreted as a branching process (4, 18), where an active neuron has a finite probability of activating its neighbors. When each active neuron induces, on average, the firing of a single neighbor, the system is thought to be in a critical state, where the activity neither explodes nor always dies out quickly. This results in a scale-free distribution of activation sequences observables, namely the size and duration of the neuronal avalanches. A key assumption in this picture is that there exists a separation of time scales, where the spontaneous activations of neurons (those that initiate an avalanche) happen much more slowly than the spreading of that avalanche across the population, i.e., there only exists a single avalanche at any given time. However, in most neuronal systems, that is far from the truth (19, 20), and many avalanches can coexist simultaneously, resulting in a more difficult interpretation of the observed statistics (18).

Until recently (10, 13), neuronal avalanches had mostly been described in systems where the network switches between periods of high and low-frequency activity such that the periods of high activity dominated the neuronal avalanche statistics. *In vivo* and in some slice preparations, such a switching behavior corresponds to the so-called “up” and “down” states, i.e., slow cortical oscillations present during slow-wave sleep (21) that are initiated by pyramidal neurons near layer V and propagate towards the other cortical layers. In other preparations, like dissociated cultures (19), the bistable behavior is more akin to hippocampal sharp wave-ripples (22), which also have a well-defined size and duration. Theoretically, this phenomenon has often been described in terms of synchronization (23) or as spontaneous switching around a bifurcation (24). However, both cortical up and down states and hippocampal short-wave ripples have a strong spatial component, initiating at specific sites and propagating throughout the tissue, reminiscent of a classical spatially-extended excitable system (25).

The specific hypothesis of spontaneous switching around a bifurcation implies that the dynamics in the high-activity state are critical whereas it is subcritical in the low-activity state (24). Here, we test this hypothesis explicitly in neuronal cultures that do show alternating activity behavior by analyzing the two different states separately. We find that both high-activity and low-activity states can exhibit similar critical signatures if appropriate time scales are chosen for defining neuronal avalanches. These results point to the existence of different mechanisms of neuronal communication, with different time scales, acting during either high-activity or low-activity states. We also show that detailed model simulations of dissociated cultures follow the aforementioned hypothesis and are, thus, incompatible with the experimental observations for the low-activity state. This suggests the existence of processes with long timescales (of the order of a few hundred ms) that play a significant role in shaping the dynamics during the low-activity state that are currently not captured by existing models.

## Materials and methods

### Hippocampal cultures

Cultures from dissociated hippocampal neurons and glial cells, prepared from newborn P0 Sprague-Dawley rats, were plated on Si chips of 1 mm thickness, and 1 cm^2^ surface area, placed on individual 24-well plate wells; as described previously (26, 27). Each chip was Matrigel-coated (Beckton Dickinson) and placed in Basal Medium Eagle (BME). Cells were initially plated at a density of 30,000 cells/ml. The culture medium was not changed during the first week and every four days thereafter.

### Calcium imaging

Cultures were grown for up to 2-3 weeks before imaging. Prior to imaging, cultures were incubated with Fluo-4 calcium indicator for 20 min. Afterward, cultures were washed and placed in an individual well with an extracellular bath solution (EBS) containing 135 mM NaCl, 10 mM glucose, 3 mM CaCl2, 5 mM KCl, 2 mM MgCl2 and 5 mM Hepes, pH was adjusted to 7.3 with NaOH, and osmolarity to 310 mOsm with Sorbital. Calcium imaging was performed following Refs. (10, 26). In brief, we used a high temporal resolution camera that allowed us to record neuronal activity at 200 fps (Hammamatsu Orca-Flash 4.0) on an upright microscope with low magnification (field of view of *L* ≈ 500 *μ*m). We recorded the spontaneous, non-stimulated fluorescence neuronal activity of up to 1500 neurons for 20 min (pixel resolution of 0.65*μ*m/pixel). With this setup we are able to record most neuronal activity within a large field of view with single-cell resolution and a temporal resolution comparable to the fastest timescales of synaptic integration; avoiding several of the drawbacks caused by spatial and temporal sub-sampling and hidden neurons (28, 29).

### Pharmacology

In a subset of experiments, we blocked inhibitory connections by performing bath application of picrotoxin (PTX), a noncompetitive GABA_A_ receptor antagonist, with a concentration of 50 *μ*M during 15 min at room temperature after the first imaging session. Cultures were then typically imaged in a different field of view for an additional 20 min.

### Data preprocessing

Data preprocessing of calcium imaging experiments was performed as previously described in (30) using the NETCAL software (31) platform (see Fig.1). In brief, cell ROIs were automatically detected using a simple thresholding procedure on the time-averaged image of the recording and posteriorly cleaned up with morphological opening operations. Time series for each ROI were extracted, detrended, and normalized to Δ*F/F*_0_ units (where *F*_0_ was computed prior to the detrending operation). Spike inference was performed using the OASIS algorithm (32). See Table 1 for a summary of the list of recording experiments and their properties.

**Fig. 1.**
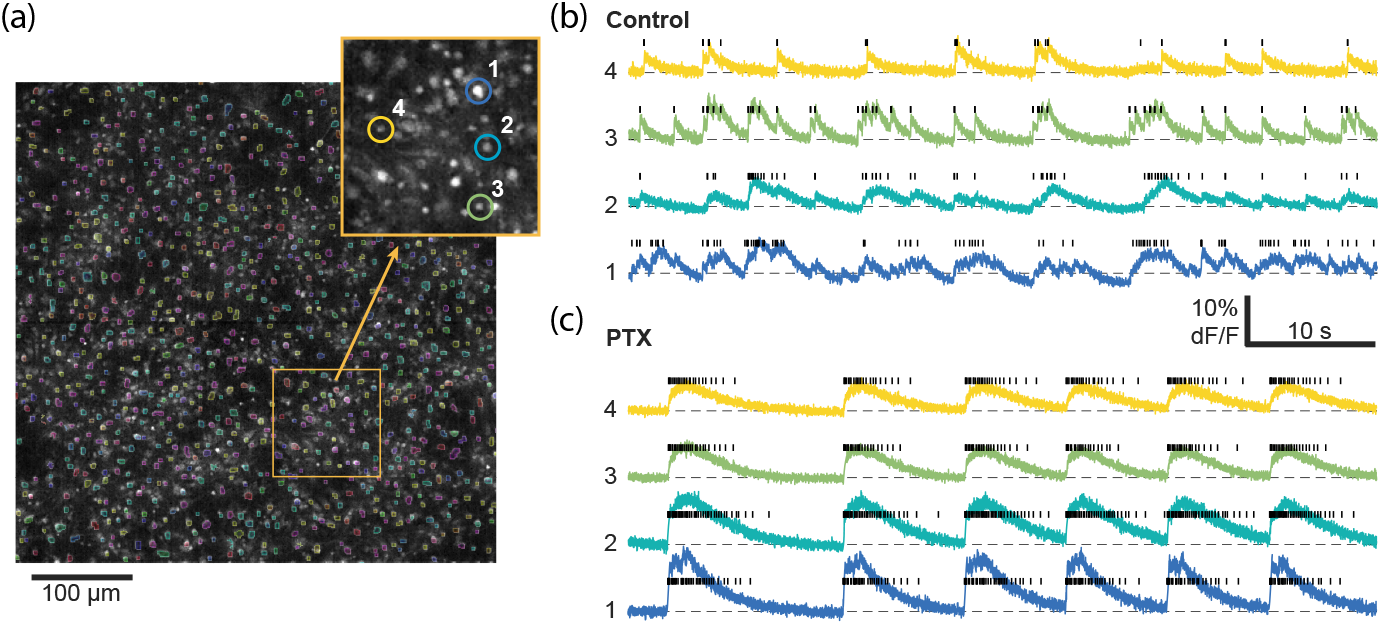
Calcium imaging analysis workflow. (a) 1000-1500 cells are automatically segmented within the field of view based on their average fluorescence. Inset: Zoom in on the field of view with 4 characteristic cells highlighted. Right: Timeseries from the same 4 highlighted cells as well as their inferred spike trains (black bars) before (b) and after (c) PTX application. For this particular example, we imaged the same cells before and after PTX. Only a smaller time window out of the 20 min recordings is shown.

**Table 1.** Summary of the properties of different recordings and simulations, including the presence (E+I) or absence (E) of active inhibitory cells as well as the average interspike interval (ISI) per neuron ⟨ISI⟩ during the up and down states.

| Recording | Type | # Cells | $\langle \text{ISI} \rangle_{\text{up}}$ (ms) | $\langle \text{ISI} \rangle_{\text{down}}$ (ms) |
| --- | --- | --- | --- | --- |
| 1 | E+I | 1374 | 216 | 2955 |
| 2 | E+I | 1510 | 211 | 1438 |
| 3 | E | 1426 | 210 | 419 |
| 4 | E+I | 825 | 86 | 449 |
| 5 | E | 813 | 407 | 1444 |
| 6 | E+I | 998 | 70 | 245 |
| 7 | E | 1001 | 248 | 613 |
| 8 | E+I | 893 | 106 | 620 |
| 9 | E | 875 | 404 | 1409 |

### Mathematical model and simulations

The dynamics of cultures from dissociated neurons were modeled and simulated following (19, 25). In brief, for the ‘homogeneous’ simulations, single neuron dynamics are modeled by a quadratic integrate and fire model with adaptation (33–35), i.e.,

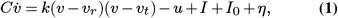

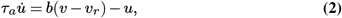

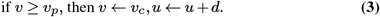

Here, Eq. (1) corresponds to the dynamics of the soma membrane potential *υ* (*t*) and *υ*_*r*_ = − 60 mV and *υ*_*t*_ = − 40 mV are the resting and threshold potentials, respectively. *C* = 100 ms is the normalized membrane capacitance and *u* models an inhibitory current that represents internal slow currents generated by the activation of ion channels. *I* accounts for the synaptic inputs from other neurons. *I*_0_ represents the spontaneous release of synaptic vesicles (minis), which are modeled as a Poisson process with frequency *λ* = 0.1 ms^*−*1^. Each mini generates a current with amplitude *g*_*m*_ = 30 mV which decays exponentially with a time constant *τ*_*m*_ = 10 ms. *η* is a white noise term with autocorrelation ⟨ *η*(*t*)*η*(*t*^′^) ⟨ = 2*g*_*s*_*δ*(*t* − *t*^′^) with *g*_*s*_ = 30 mV. Eq. (2) is the evolution of the slow currents and *τ*_*a*_ = 33 ms, *k* = 0.7 mV^*−*1^, *b* = − 2 and *d* = 100 mV are parameters that control recovery and adaptation. Every time a given neuron *i* fires, it produces an (excitatory or inhibitory) current on its output neighbors of the form

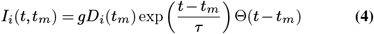

where *t*_*m*_ is the spike time and *g* is the synaptic strength. Its value for excitatory synapses is used as a control parameter to obtain the desired time interval separating subsequent up states (which is achieved for *g* ≈ 40 mV) while the inhibitory strength is fixed at − 50 mV. *τ* is the characteristic time constant (10 ms for excitatory currents and 20 ms for inhibitory ones). Θ(*t*) is the Heaviside function and *D* short-term synaptic depression (STD). The evolution of STD is described by

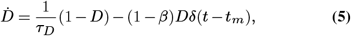

where *τ*_*D*_ (2 s for excitatory and 0.2 s for inhibitory cells) is the synaptic recovery time, and *β* (0.8 for excitatory and 0.95 for inhibitory cells) controls the level of depression after each spike.

To mimic the experimental cultures and establish realistic connectivity patterns, neurons were placed randomly on a square region with 10 mm sidelength until a density of *ρ* = 800 neurons/mm^2^ was reached (80,000 neurons total). For each neuron, an axon was grown as a biased random walk with total length given by a Rayleigh distribution with variance 900*μ*m^2^. Starting from the cell body, a starting angular direction was picked randomly, and a segment of 10*μ*m was grown. At the end of the segment, a new direction was chosen, centered on the previous one, following a Gaussian distribution with a standard deviation of 15°. This process was repeated until the desired total length was reached. For each neuron, a dendritic tree was modeled as an effective circular interaction area with a radius obtained from a Gaussian distribution with mean 150*μ*m and standard deviation 40*μ*m. If an axon crossed the dendritic tree of another neuron, a connection was established with probability 0.13. Finally, 20% of the neurons were randomly chosen to be inhibitory and the remaining ones excitatory.

For the ‘heterogeneous’ simulations we lowered the standard excitatory population to 70% and added 10% of excitatory bursty cells (accomplished by changing *υ*_*c*_ = − 40 and *d* = 50). To all excitatory cells, we changed the resting membrane potential to a base value of *υ*_*r*_ = − 62 mV and added an offset drawn from a Rayleigh distribution with standard deviation *σ* = 2 mV, to add variability to the firing rates consistent with experimental data.

Each simulation had a fixed run length of 1 h and was simulated with a first-order Euler algorithm with a time step of 0.1 ms. Random numbers were generated with the MTGP32 implementation of the Mersenne Twister for the GPU (36) and initialized with a random seed for each simulation.

### Detection of up and down states

The detection of up and down states is done based on thresholding the population firing rate. The firing rate for each frame is defined as the number of spikes in that frame normalized by the number of neurons. A Gaussian kernel with *σ* = 5 frames for experimentally recorded data and *σ* = 20 frames for simulated data is used to smooth the firing rate traces. To separate up and down states we used a Schmitt trigger (37), thresholding the normalized activity with an upper threshold = 0.001 and a lower threshold = 0.0003. We kept the same criteria for all recordings and simulations.

### Neuronal avalanches and scaling collapse procedure

Following the standard approach (4), a neuronal avalanche is defined as the largest sequence of consecutive time bins containing spikes in every single time bin, separated by time bins during which none of the neurons in the culture fire. The avalanche duration, *T*, corresponds to the number of time bins, and the avalanche size, *S*, is the total number of spikes over the duration of an avalanche. Based on this definition, for a fixed number of neurons, it is expected that the choice of the size of the time bin affects the avalanche statistics, specifically the size *S* and duration *T* of avalanches. To capture this, we follow here the standard finite size scaling approach used in the context of phase transitions (see, for example, (38)). In our context, its original formulation in terms of a varying number of neurons can be recast in terms of a varying size of the time bin. It is based on the hypothesis that for a fixed number of neurons, the effect of temporal bin size on scale-free avalanche statistics (e.g., avalanche size distribution, *P* (*S*)) can be taken into account as follows:

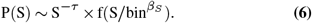

This functional form indicates, for suitably chosen scaling exponents *τ* and *β*_*S*_, the distributions for different bin sizes can be collapsed onto the scaling function *f* by plotting 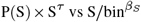. In the case of exponential-like avalanche statistics (*τ* = 0), one can achieve a similar data collapse with a scaling exponent *β*_*S*_ when plotting 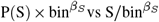.

To replicate the experimental field of view (which covers only a small fraction of the whole culture) in the simulations, we only analyzed the avalanche statistics of neurons within a circular patch of radius *r* = 0.5 mm for the E simulations and *r* = 0.8 mm in the E+I simulations. This radius was chosen to ensure that the total activity rate in the up states was of the order of 1 event per frame or time step (0.1 ms). Each of these patches contained in total 400 to 1000 neurons. To increase the number of avalanches and obtain reliable estimates of the avalanche statistics, for each simulation and each experiment, we randomly selected 100 different local patches consisting of 50% of the neurons and combined the avalanches from all patches into a single distribution. This approach also allows us to obtain avalanche statistics for different temporal bin sizes.

### P-value estimation for power-law distributions

After identifying the optimal exponents *α* and *β*, using the scaling collapse procedure (Eq. (6)), our next step is to assess the validity of the power-law model as a hypothesis for the specific recording or simulation. To achieve this, we check whether we can identify an extended range for the size or duration in the avalanche distribution function that gives a reasonably high p-value (*>* 0.1). A detailed description of the process of identifying the range is presented in the captions of Fig. S1. To find the pvalue, we first used the Kolmogorov-Smirnov (KS) statistic. KS statistic quantifies the distance between two probability distributions (39), which is defined as the maximum distance between the cumulative distribution functions (CDFs) of the two distributions (here data and fitted model):

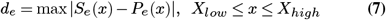

Here, *S*_*e*_(*x*) is the CDF of the empirical data, and *P*_*e*_(*x*) is the CDF corresponding to the fitted model. The fitted model is estimated using the power-law scaling procedure related to Eq. 6. The reported p-value is the probability of observing a KS value bigger than *d*_*e*_ for synthetic data generated by the fitted model and provides a measure of whether it is likely that the empirical data do indeed follow the fitted model. One can show that its value can be calculated from the following theoretical expression (40):

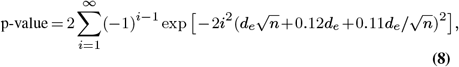

where *n* is the number of samples in the data set. As mentioned above, to enhance our statistics we identified avalanches over 100 iterations (for both experimental and simulation data). The reported p-value is calculated over each of those iterations. The reported p-value is the mean ± standard error of the mean (SEM), where the mean value and SEM are calculated over 100 iterations. See Fig. S1 for a representative example of the procedure and Table S1 for fitting details.

## Results

### Neuronal cultures

The overall activity of two to three-week-old neuronal cultures typically switches between high activity periods or ‘up states’ and low activity periods or ‘down states’ as shown in Fig. 2. The up states — often also referred to as network bursts — occur quasiperiodically and involve the vast majority of all neurons. As Fig. 2 also shows, their duration is quite regular as well. The activity during up states can be understood as a set of causal cascades of induced firings across the observed population of neurons in the culture, often modeled as a branching process (4). The cascade of neuronal activity is studied in the framework of neuronal avalanches as described in the *Materials and Methods* section. Previous studies have found that neuronal avalanches exhibit statistics of a branching process at or close to its critical point (4, 41). In the often-observed case of a mean-field branching process, the activity propagates on a tree-like network without feed-back loops and the avalanches follow a scale-free behavior, i.e.:

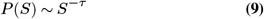

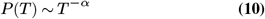

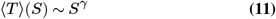

where *P* is the probability distribution function (PDF) of the associated variable and *τ* = 1.5, *α* = 2.0, and *γ* = 0.5 are the critical meanfield exponents (41). These critical exponents — whether they take on mean-field values or not — are necessarily related through the scaling relation:

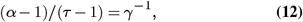

which provides an additional test for the presence of critical behavior. Here, we study the avalanche statistics of the up state and the down state separately. Due to the vastly different activity levels, the corresponding ISIs used to define the neuronal avalanches are also vastly different. For our experimentally recorded data the single cell ISIs are: ISI_up state_ = 217 ms ± 43 ms and ISI_down state_ = 1065 ms ± 302 ms (mean ± SEM). Nevertheless, we find similar statistical features across up and down states. Examples of distributions of avalanche sizes and durations as well as the relation between sizes and durations are plotted in Fig. 3. Indeed, for both the up and down state in Fig. 3, we recover Eqs. (9-11) and the exponents are close to the mean-field values. As Table 2 shows, this is true for the up state even if we block all inhibitory connections by the application of a saturating concentration (50*μ*M) of picrotoxin (see *Materials and Methods* for details). Overall, we find that the majority of experimental recordings have an up state that is consistent with power-law statistics in their sizes over an extended range (see Table S1). Such power-law behavior is slightly less prevalent in the down state (Tables 2, S1), but still prominent. Moreover, in all cases the scaling relation Eq. (12) holds within the statistical uncertainties (Fig. S2), consistent with critical behavior.

**Fig. 2.**
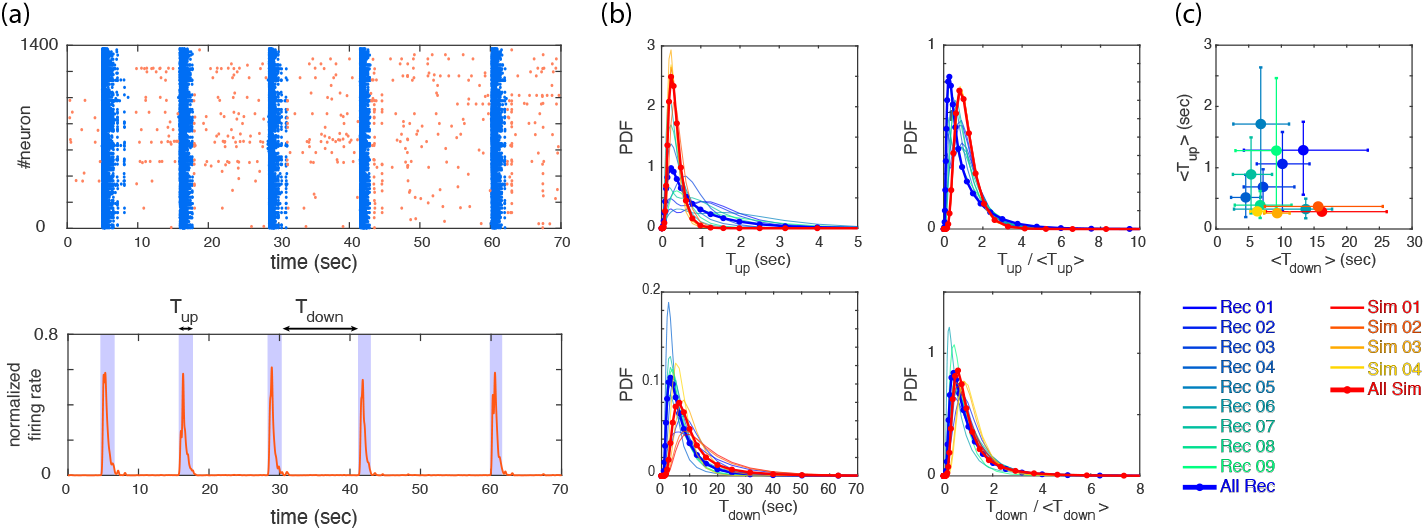
(a) Example of a raster plot for an experimental recording (top panel) and its corresponding normalized firing rate (bottom panel) are depicted. The up and down states are identified by thresholding the firing rate, where bins of data with firing rate*>*threshold are identified as up state and bins of data with firing rate less threshold are identified as down state (see *Materials and Methods* for details). We define *T*_up_ and *T*_down_ to measure the lengths of up and down states as visualized in the bottom panel. (b) Probability Density Functions (PDFs) of *T*_up_, *T*_down_, *T*_up_ */ < T*_up_ *>*, and *T*_down_ */ < T*_down_ *>* are shown, where each curve represents one of the experimental recordings or simulations (see legends). (c) The mean values of *T*_up_ and *T*_down_ for all the recordings and simulations are shown. Error bars correspond to 75 percentiles and the same color coding as panel (b) is used.

**Table 2.**
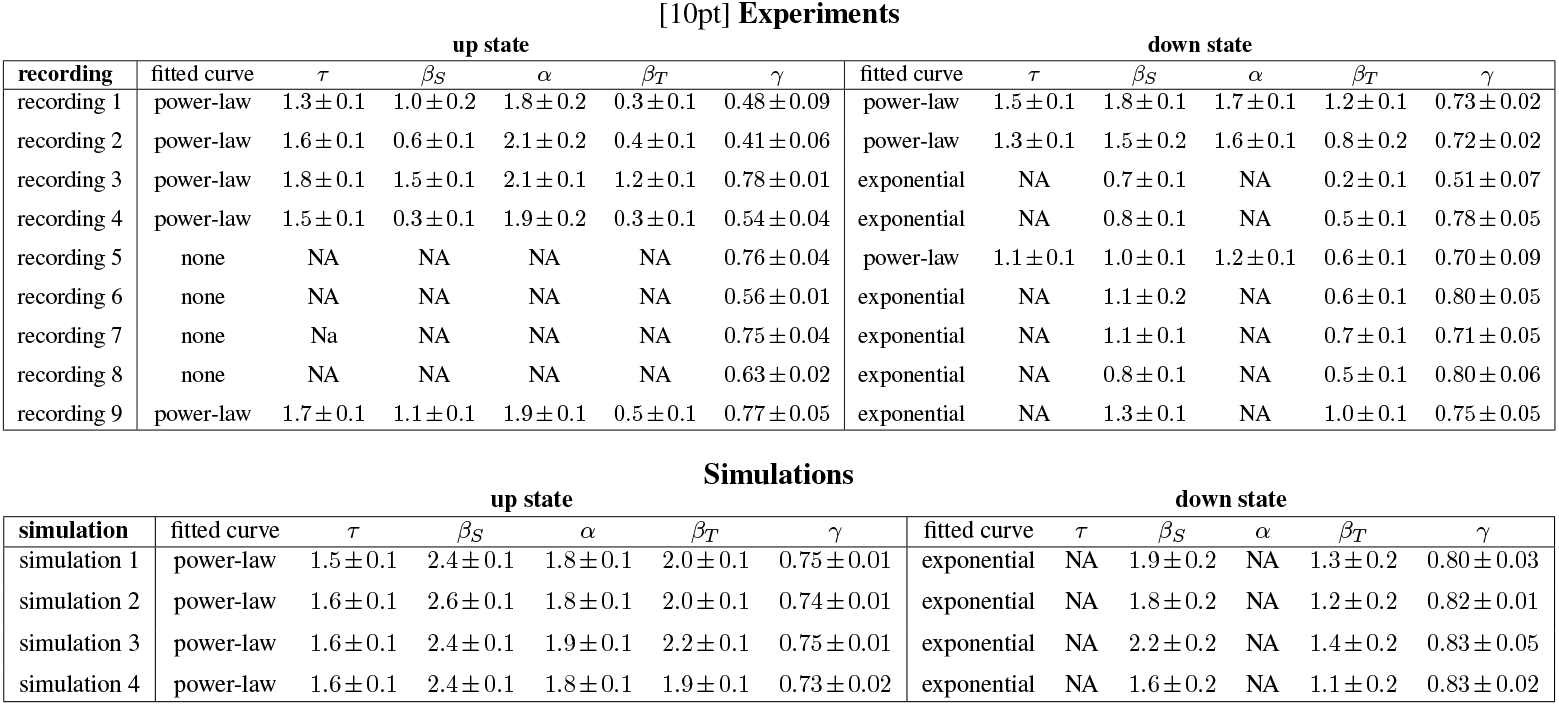
This table summarizes the exponents for all of the recordings and simulations. *τ, α* and *β*_*S*_ and *β*_*T*_ are calculated using the scaling procedure described in *Materials and Methods*. In some cases, the power-law behaviour was too limited in range (less than one decade) to reliably estimate the exponents, explaining the absence of a fitted curve. *γ* is calculated using least-squares fitting on the log of avalanche sizes and durations, where we use 95% confidence bounds to estimate the error.

[10pt] Experiments
| recording | up state |  |  |  |  |  | down state |  |  |  |  |  |
| --- | --- | --- | --- | --- | --- | --- | --- | --- | --- | --- | --- | --- |
| | fitted curve | $\tau$ | $\beta_S$ | $\alpha$ | $\beta_T$ | $\gamma$ | fitted curve | $\tau$ | $\beta_S$ | $\alpha$ | $\beta_T$ | $\gamma$ |
| recording 1 | power-law | $1.3 \pm 0.1$ | $1.0 \pm 0.2$ | $1.8 \pm 0.2$ | $0.3 \pm 0.1$ | $0.48 \pm 0.09$ | power-law | $1.5 \pm 0.1$ | $1.8 \pm 0.1$ | $1.7 \pm 0.1$ | $1.2 \pm 0.1$ | $0.73 \pm 0.02$ |
| recording 2 | power-law | $1.6 \pm 0.1$ | $0.6 \pm 0.1$ | $2.1 \pm 0.2$ | $0.4 \pm 0.1$ | $0.41 \pm 0.06$ | power-law | $1.3 \pm 0.1$ | $1.5 \pm 0.2$ | $1.6 \pm 0.1$ | $0.8 \pm 0.2$ | $0.72 \pm 0.02$ |
| recording 3 | power-law | $1.8 \pm 0.1$ | $1.5 \pm 0.1$ | $2.1 \pm 0.1$ | $1.2 \pm 0.1$ | $0.78 \pm 0.01$ | exponential | NA | $0.7 \pm 0.1$ | NA | $0.2 \pm 0.1$ | $0.51 \pm 0.07$ |
| recording 4 | power-law | $1.5 \pm 0.1$ | $0.3 \pm 0.1$ | $1.9 \pm 0.2$ | $0.3 \pm 0.1$ | $0.54 \pm 0.04$ | exponential | NA | $0.8 \pm 0.1$ | NA | $0.5 \pm 0.1$ | $0.78 \pm 0.05$ |
| recording 5 | none | NA | NA | NA | NA | $0.76 \pm 0.04$ | power-law | $1.1 \pm 0.1$ | $1.0 \pm 0.1$ | $1.2 \pm 0.1$ | $0.6 \pm 0.1$ | $0.70 \pm 0.09$ |
| recording 6 | none | NA | NA | NA | NA | $0.56 \pm 0.01$ | exponential | NA | $1.1 \pm 0.2$ | NA | $0.6 \pm 0.1$ | $0.80 \pm 0.05$ |
| recording 7 | none | NA | NA | NA | NA | $0.75 \pm 0.04$ | exponential | NA | $1.1 \pm 0.1$ | NA | $0.7 \pm 0.1$ | $0.71 \pm 0.05$ |
| recording 8 | none | NA | NA | NA | NA | $0.63 \pm 0.02$ | exponential | NA | $0.8 \pm 0.1$ | NA | $0.5 \pm 0.1$ | $0.80 \pm 0.06$ |
| recording 9 | power-law | $1.7 \pm 0.1$ | $1.1 \pm 0.1$ | $1.9 \pm 0.1$ | $0.5 \pm 0.1$ | $0.77 \pm 0.05$ | exponential | NA | $1.3 \pm 0.1$ | NA | $1.0 \pm 0.1$ | $0.75 \pm 0.05$ |

**Table 2.** This table summarizes the exponents for all of the recordings and simulations.
| simulation | up state |  |  |  |  |  | down state |  |  |  |  |  |
| --- | --- | --- | --- | --- | --- | --- | --- | --- | --- | --- | --- | --- |
| | fitted curve | $\tau$ | $\beta_S$ | $\alpha$ | $\beta_T$ | $\gamma$ | fitted curve | $\tau$ | $\beta_S$ | $\alpha$ | $\beta_T$ | $\gamma$ |
| simulation 1 | power-law | $1.5 \pm 0.1$ | $2.4 \pm 0.1$ | $1.8 \pm 0.1$ | $2.0 \pm 0.1$ | $0.75 \pm 0.01$ | exponential | NA | $1.9 \pm 0.2$ | NA | $1.3 \pm 0.2$ | $0.80 \pm 0.03$ |
| simulation 2 | power-law | $1.6 \pm 0.1$ | $2.6 \pm 0.1$ | $1.8 \pm 0.1$ | $2.0 \pm 0.1$ | $0.74 \pm 0.01$ | exponential | NA | $1.8 \pm 0.2$ | NA | $1.2 \pm 0.2$ | $0.82 \pm 0.01$ |
| simulation 3 | power-law | $1.6 \pm 0.1$ | $2.4 \pm 0.1$ | $1.9 \pm 0.1$ | $2.2 \pm 0.1$ | $0.75 \pm 0.01$ | exponential | NA | $2.2 \pm 0.2$ | NA | $1.4 \pm 0.2$ | $0.83 \pm 0.05$ |
| simulation 4 | power-law | $1.6 \pm 0.1$ | $2.4 \pm 0.1$ | $1.8 \pm 0.1$ | $1.9 \pm 0.1$ | $0.73 \pm 0.02$ | exponential | NA | $1.6 \pm 0.2$ | NA | $1.1 \pm 0.2$ | $0.83 \pm 0.02$ |

Our analysis of the experimental data shows in particular that the statistical behavior of the neuronal avalanches in the down state can be statistically indistinguishable from the up state, especially if one focuses on *τ*, see Fig. 3 and Table 2. Note that the different ranges in duration and sizes (see also Table S1) are due to the shorter relative duration of the down states with respect to their corresponding ISI, as a comparison with the case of experimental recordings with continuously low steady-state activity (which can be interpreted as a system without an up state and instead being exclusively in a down state) (10) confirms.

**Fig. 3.**
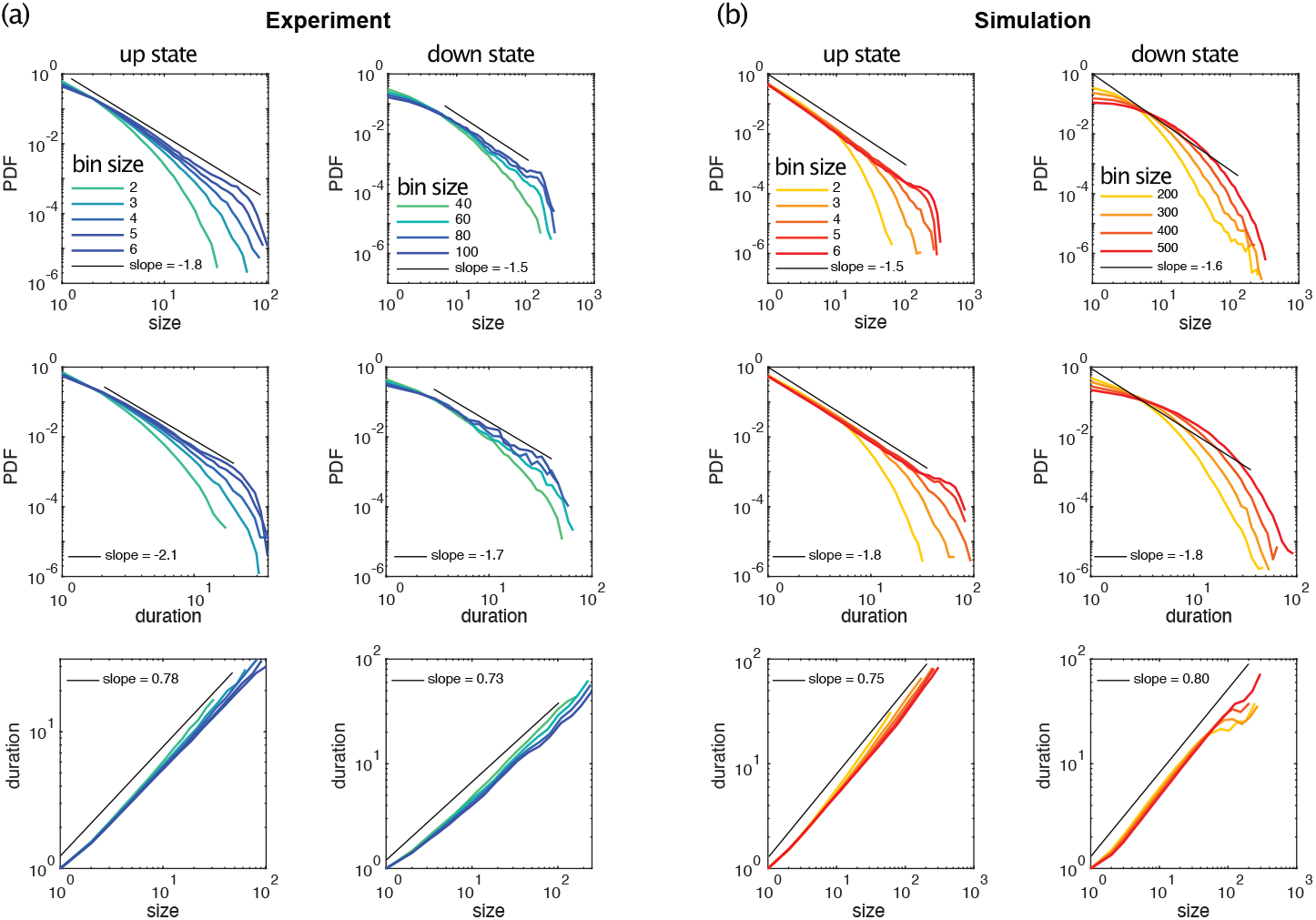
Example distributions of neuronal avalanche sizes and durations for up and down states of experimental data and simulated data are plotted for different temporal bin sizes. **(a)** The up state is from experimental recording 3, the down state is from recording 1. **(b)** Simulation 1 is shown but all simulations exhibit almost identical avalanche statistics.

As outlined in the *Materials and Methods* section, in this study we take advantage of a scaling analysis that also yields critical exponents denoted as *β*_*S*_ and *β*_*T*_ that can capture finite size behavior. This provides us with an additional tool for characterizing the dynamics of neuronal systems within the framework of neuronal avalanche statistics. As depicted in Fig. 4(a, b), we studied the scaling properties of two types of distributions: i) Power-law scaling which gives us *β*_*S*_ (*β*_*T*_ ) and *τ* (*α*), and ii) Exponential-like scaling that gives us only *β*_*S*_ (*β*_*T*_ ), for avalanche sizes (durations). The estimate of the exponents becomes more reliable when the scaling collapse is obtained for a wider range of varying bin sizes. The summary of all estimated exponents for i), along with the corresponding p-values for the reported critical exponents *τ* and *α*, (see *Materials and Methods* for details) for both up and down states, are plotted in Fig. 4(c). While the variation in *τ* and *α* is rather small, especially in the up state, this is not the case for *β*_*S*_ and, to a lesser degree, for *β*_*T*_ . This higher variability is also visible in Fig. 4(d), which displays the corresponding mean values and uncertainties for all exponents for both the up and down states. Fig. 4(e) displays the mean values of *β*_*S*_ and *β*_*T*_ for distributions exhibiting exponentiallike scaling ii), which exclusively occurs in the down state, see Table 2.

**Fig. 4.**
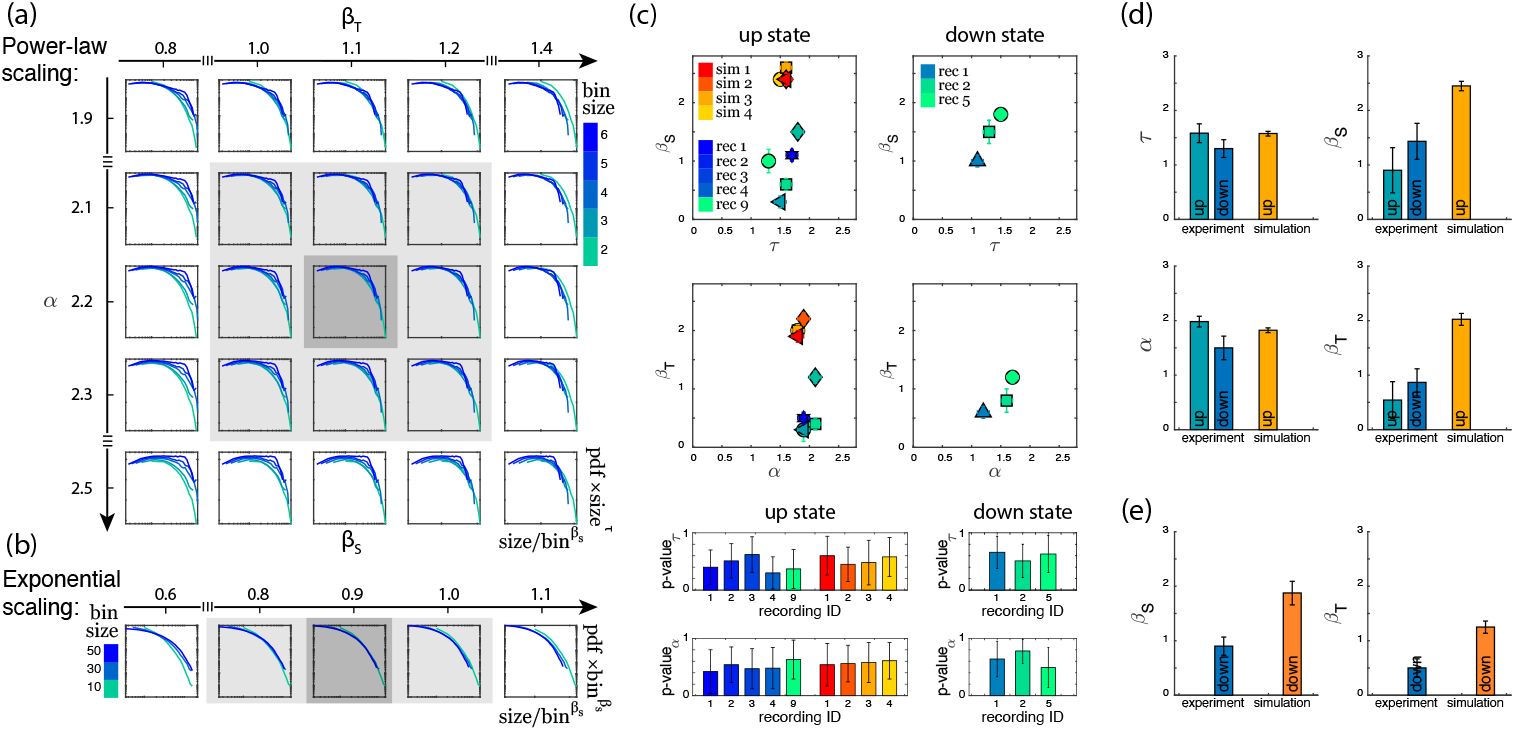
(a) The power-law scaling collapse procedure for finding the exponents is visualized for an example experimental dataset (recording 3, up state, avalanche duration). For details of this procedure, see *Materials and Methods*. (b) The exponential-like scaling collapse procedure for finding the scaling exponent *β* for a sample experimental data is visualized (recording 3, down state, avalanche sizes). (c) Summary of the exponents for size and duration of all recordings during up and down states are presented here. Each point represents an experimental or a simulated dataset. We have only included recordings that exhibit power-law behavior over extended ranges as indicated in Table 2. P-values of the fitted power laws with exponents *τ* and *α*, respectively, are shown in the bottom panels (mean + std of the ensemble of subpopulations). For each of the two columns, the same color scheme as the top panel is preserved. (d) Average of all power-law exponents for the data sets in (c). (e) Average of *β*_*S*_ and *β*_*T*_ for all data sets that show exponential-like behavior as the example in (b).

### Modeling results

The similarity between the up state and the down state in terms of the neuronal avalanches for our experimental recordings suggests that either (a) there are different mechanisms of information processing in this neuronal system with different associated time scales, or (b) the concept of neuronal avalanches is not specific enough to establish insight into the differences between up and down states. To investigate this further, we employed several computational models that try to mimic the behavior and dynamics of neuronal cultures as described in *Materials and Methods*.

In cultures from dissociated neurons, a down-to-up state transition occurs stochastically, due to a competition between noise-induced activations and recovery from synaptic depression (19). When the two timescales (activations and recovery) are very different, one dominates, and the behavior might appear quasi-periodic, but is still noise-dominated. On the other hand, transitions from up to down state are mediated primarily by the depletion of neurotransmitters caused by the high-frequency firings during the up state (42, 43). Noise plays a much lesser role in this transition, and the relative variance in the duration of the up states is much narrower than in the time interval separating subsequent up states. The models used here are able to reproduce most of these macroscopic observables as Fig. 2 shows. However, when applying the same methodology to characterize the avalanche statistics as in the experimental data, there are important differences. In the simulations, the avalanches observed during the up states followed powerlaw distributions with slopes of *τ* ≃− 1.5 and *α* − 1.8 for sizes and durations, respectively, see Fig. 3b left and Table 2. These avalanche statistics also satisfied the scaling relation Eq. (12) within the statistical uncertainties (Fig. S2). During the down state, however, the avalanche statistics were always far from a power-law distribution, suggesting a sub-critical or exponential-like behavior instead (see Fig. 3b right). As a result, the down state in the simulated data is absent from Fig. 4(d).

Although the scaling exponents *τ* and *α* were largely consistent during the up states with those observed experimentally (see Table 2), the scaling exponents *β*_*s*_ and *β*_*T*_ differed substantially from the experimental ones as follows from Fig. 4(d) and (e). The measured exponents across the different simulation conditions were robust, with little variability. The presence or absence of inhibitory connections, as well as heterogeneous cell populations (see *Materials and Methods*), produced no significant differences in the avalanche statistics. Changes in connectivity strength, synaptic depression parameters (depression strength and recovery time constant), and the presence of other synaptic currents (NMDA), produced no changes in the reported exponents either (not shown). Similarly, although the metric properties of the network structure have a large impact on the presence of the up-down transition (44), changes in connectivity, i.e., between metric-embedded to a random graph, also resulted in no changes in any of the exponents.

## Discussion

In this work, we present for the first time the simultaneous characterization of neuronal avalanches in dissociated neuronal cultures during both up and down states. In several cultures, the avalanche statistics during both up and down states show critical behavior, as indicated by avalanche sizes and durations. In addition, the associated critical exponents satisfy the corresponding scaling relation (Eq. (12)), supporting the presence of critical behavior. These exponents are largely within the range of those of a critical mean-field branching process, in line with those previously reported across many preparations (4, 10, 41). The exponents associated with finite-size scaling show some differences between up and down states. Yet, the variations are large and the statistics is rather limited such that it is difficult to make any definite statements. More importantly, we find that the scaling relation *β*_*T*_ = *γβ*_*S*_ (which follows directly from Eq. (11)) holds within the statistical uncertainties in almost all cases (Fig. S2), providing further evidence of criticality. The mechanisms by which these cultures can self-organize to maintain critical avalanche statistics across very different activity regimes are still unknown. If differences in the exponents were indeed present, this would further indicate that the two activity regimes are associated with distinct universality classes.

There has been extensive theoretical and modeling work to try and describe the up and down states as an emergent property of neuronal systems. However, only a few models can simultaneously reproduce the experimentally observed avalanche statistics and switch between up and down states. These include integrate-and-fire networks with structured connectivity through learning (45); networks with scale-free connectivity coupled to a global field for the up and down switching (46); self-organized criticality around a saddle-node bifurcation (24); and a mesoscopic model on the verge of a synchronization transition (47). However, only Ref. (24) treated the avalanche statistics of up and down states separately. They introduced a self-organized critical model with noise-driven, spontaneous transitions between up and down states. The transitions occur around a bifurcation such that the down state is subcritical and the up state is critical. Such a framework, however, is not applicable to our experimental system since its predictions are inconsistent with several experimental observations. Namely, (i) up states possessing characteristic durations, with a well-defined mean and variance (see, for example, Fig. 2); (ii) the down states presenting a welldefined duration (usually called the interburst interval, IBI) that is correlated with the slow timescale of synaptic depression (43, 48); and (iii) the activity during the down states being possibly critical, as reported here and in (10). Despite trying to take all of this into account, our computational model was still unable to reproduce some of the (new) experimental observations. Although the model was able to successfully capture many of the features of spontaneous activity in neuronal cultures (19), e.g., up and down transition statistics, distribution of states durations and their temporal correlations, up state exponents, etc., it could neither reproduce critical behavior observed during down states nor the values of the finite-size scaling exponents associated with the up states. For the model to simultaneously reproduce critical avalanche statistics during up and down states, its phase diagram would need to have critical points at two very different levels of average activity. The mechanism by which the model would be able to capture this is still unknown.

Conceptually, the fact that these cultures can dynamically transition between different levels of activity and still remain critical (or critical-like), suggests that we might have to move away from the traditional picture of a critical mean-field branching process, and even one of selforganization around a single critical point (49). Since the observed avalanche statistics need to be defined using substantially different time windows between the up and down states (based on the global interspike intervals), a model that can dynamically adapt the timescales of synaptic integration (to maintain a constant effective probability of inducing firings from a neuron to their neighbors) could be a good candidate. Glial cells are known to modulate synaptic plasticity across different timescales (50) and have recently been shown to be involved in neural communication across long timescales (51), hence being a likely candidate to support multi-scale critical dynamics. Investigating this hypothesis remains an exciting challenge for the future.

## Author Contributions

M.Y. performed the neuronal avalanche analysis guided by J.D. and J.G.O. M.Y., J.G.O. and J.D. contributed to the writing of the manuscript. M.Y. and J.G.O. performed the experimental recordings. J.G.O. performed the numerical model simulations. M.A.C. assisted with the preparation and the optical recording of cultures, and the editing of the manuscript.

## Acknowledgments

We thank Y. Hernandez and A. Kipp for their help with culture preparation and maintenance.

## Supplementary Material

**Table S1.** Power law fits - ranges and p-values

| experiment - up state |  |  |  |  |
| --- | --- | --- | --- | --- |
|  | avalanche size range | p-value | avalanche duration range | p-value |
| 1 | [4, 40] | $0.40 \pm 0.30$ | [1, 4] | $0.42 \pm 0.38$ |
| 2 | [1, 100] | $0.51 \pm 0.30$ | [1, 4] | $0.54 \pm 0.31$ |
| 3 | [3, 30] | $0.62 \pm 0.31$ | [3, 20] | $0.66 \pm 0.35$ |
| 4 | [3, 50] | $0.30 \pm 0.28$ | [1, 5] | $0.48 \pm 0.36$ |
| 9 | [2, 40] | $0.37 \pm 0.34$ | [3, 20] | $0.63 \pm 0.34$ |

| experiment - down state |  |  |  |  |
| --- | --- | --- | --- | --- |
|  | avalanche size range | p-value | avalanche duration range | p-value |
| 1 | [10, 100] | $0.66 \pm 0.28$ | [3, 30] | $0.64 \pm 0.31$ |
| 2 | [5, 60] | $0.66 \pm 0.29$ | [3, 11] | $0.78 \pm 0.26$ |
| 5 | [3, 30] | $0.44 \pm 0.32$ | [2, 12] | $0.49 \pm 0.35$ |

| simulation - up state |  |  |  |  |
| --- | --- | --- | --- | --- |
|  | avalanche size range | p-value | avalanche duration range | p-value |
| 1 | [2, 200] | $0.45 \pm 0.30$ | [2, 50] | $0.56 \pm 0.32$ |
| 2 | [2, 200] | $0.60 \pm 0.34$ | [1, 40] | $0.54 \pm 0.37$ |
| 3 | [2, 200] | $0.58 \pm 0.34$ | [1, 100] | $0.61 \pm 0.32$ |
| 4 | [2, 200] | $0.48 \pm 0.39$ | [1, 50] | $0.58 \pm 0.35$ |

**Fig. S1.**
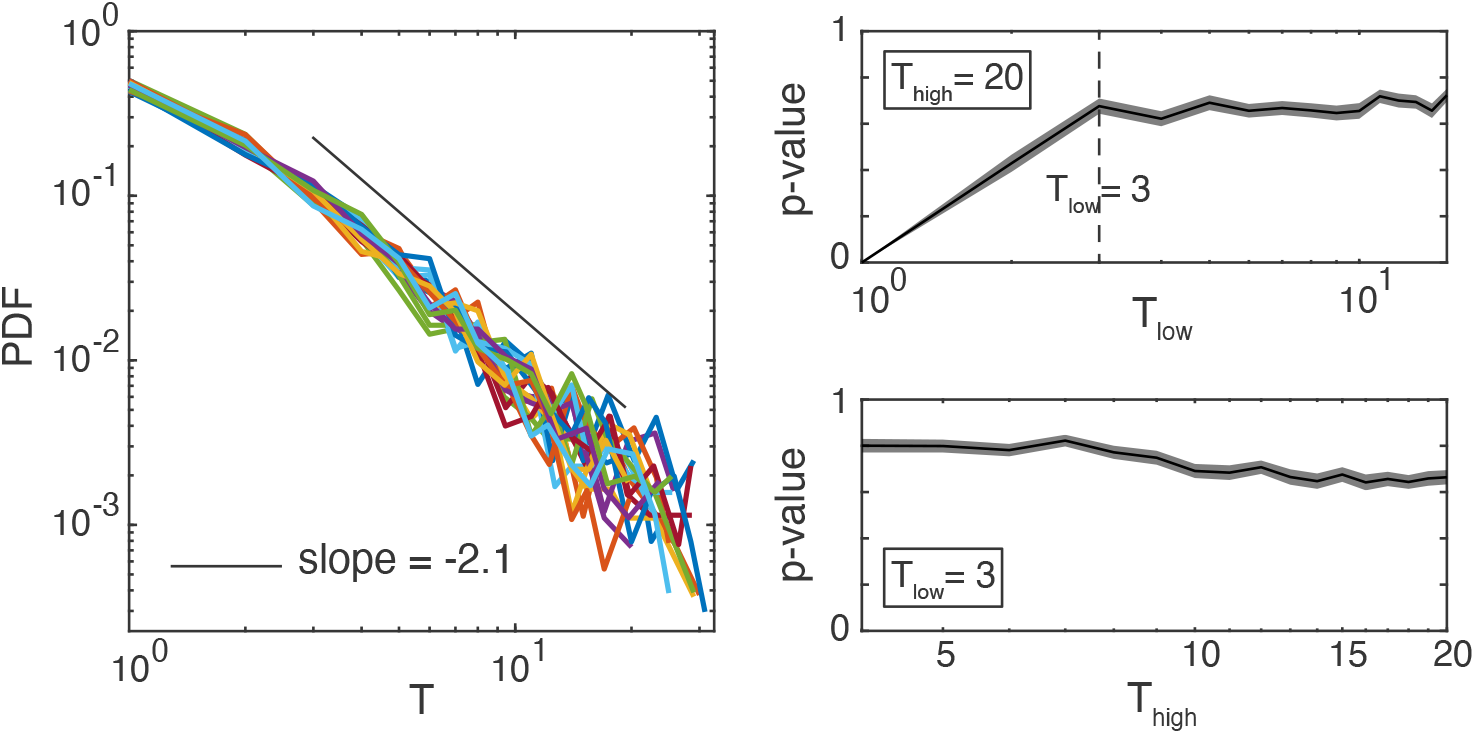
The procedure for estimating the p-value and determining the range of the power-law fit is as follows, using an example experimental recording: In the left panel, we display the probability distribution of the avalanche durations for 20 randomly selected subpopulations of neurons. The black line superimposed on this panel represents the curve fitted using the scaling procedure explained in the main text. To identify the optimal range with the highest p-value, we follow these steps: As depicted in the top-right panel, we start by manually selecting an upper bound (*T*_*high*_) and then assess the p-value while varying the lower bound (*T*_*low*_). The p-value is calculated using KS statistics as in Equations (7) and (8) from the main manuscript. Notably, we observe that *T*_*low*_ = 3 marks the point at which the p-value stabilizes at a reasonably high value. Subsequently, as illustrated in the bottom-right panel, we fix the lower bound at the previously determined value and vary the upper bound to identify the largest range that maintains a high p-value. In this particular case, the upper bound can extend to the maximum allowable value, approximately 20. Therefore, the resulting range is [3, 20], with a corresponding p-value of 0.66. In both right panels, the black line represents the mean, and the shaded area indicates the standard error of the mean (SEM) across 100 iterations.

**Fig. S2.**
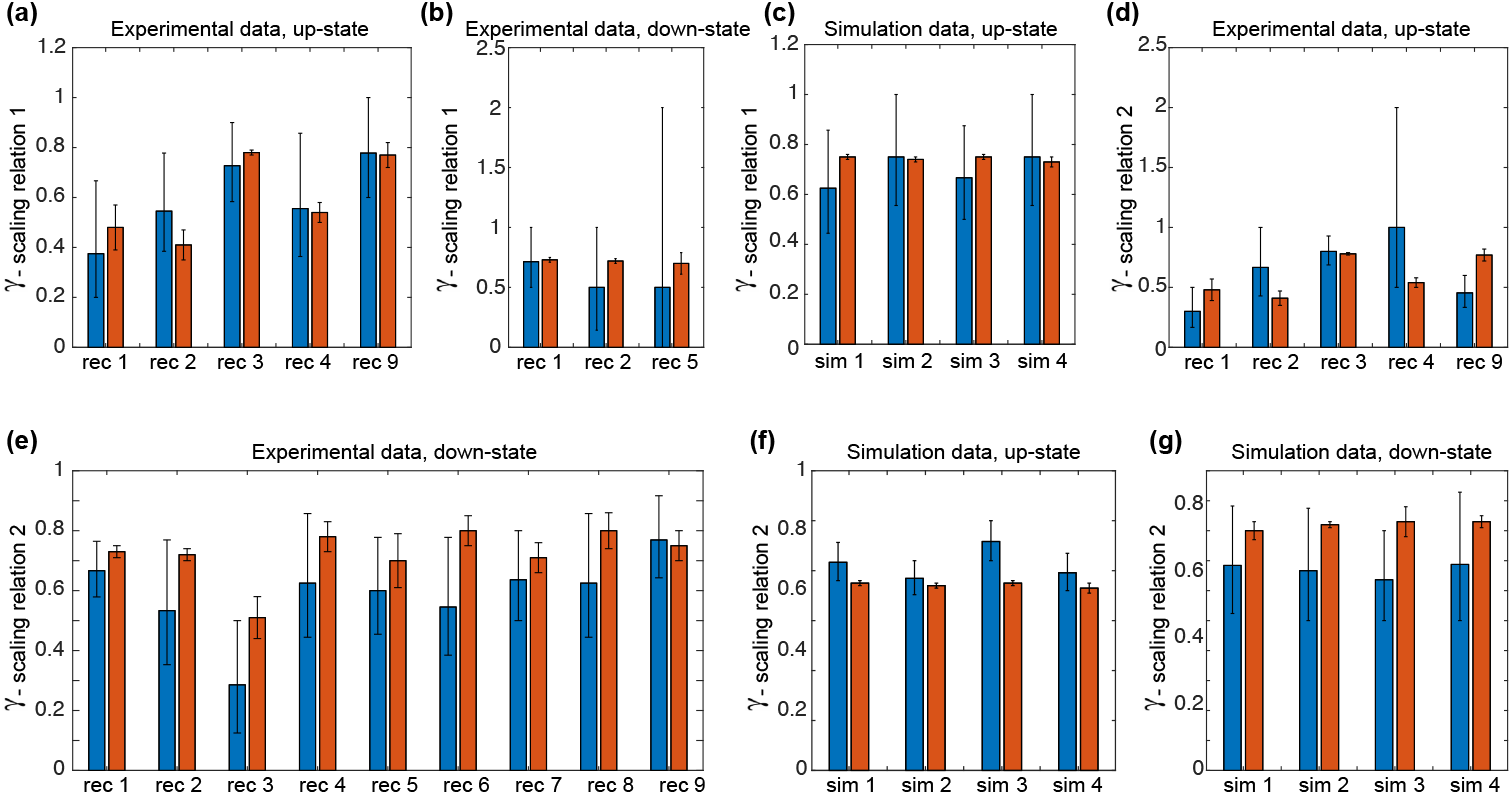
(a), (b), and (c) test the first scaling relation: *γ* = (*τ −* 1)*/*(*α −* 1), for experimental down and up states, and simulation up state data, respectively. Blue bars correspond to the predicted values of *γ* using the scaling relation (the error bars arise from the uncertaintites in *α* and *τ* ), and orange bars correspond to the directly measured values of *γ*, see Table 2 in the paper. Both values agree within the statistical uncertainties in all cases. Similarly, panels (d), (e), (f), and (g) test the second scaling relation: *γ* = *β*_*T*_ */β*_*S*_, for experimental up and down states, and simulation up and down states, respectively.

